# Decoding single-cell multiomics: scMaui - A deep learning framework for uncovering cellular heterogeneity in presence of batch Effects and missing data

**DOI:** 10.1101/2023.01.18.524506

**Authors:** Yunhee Jeong, Jonathan Ronen, Wolfgang Kopp, Pavlo Lutsik, Altuna Akalin

## Abstract

The recent advances in high-throughput single-cell sequencing has significantly required computational models which can address the high complexity of single-cell multiomics data. Meticulous single-cell multiomics integration models are required to avoid biases towards a specific modality and overcome the sparsity. Batch effects obfuscating biological signals must also be taken into account. Here, we introduce a new single-cell multiomics integration model, Single-cell Multiomics Autoencoder Integration (scMaui) based on stacked variational encoders and adversarial learning. scMaui reduces the dimensionality of integrated data modalities to a latent space which outlines cellular heterogeneity. It can handle multiple batch effects independently accepting both discrete and continuous values, as well as provides varied reconstruction loss functions to cover various assays and preprocessing pipelines. We show that scMaui accomplishes superior performance in many tasks compared to other methods. Further downstream analyses also demonstrate its potential in identifying relations between assays and discovering hidden subpopulations.

## Introduction

Recent progress in high-throughput sequencing technology made it possible to profile different omics modalities from the same cells^1^, so-called single-cell multiomics. For instance, CITE-seq offers a joint profile of transcriptomes and proteomes^2^, while single-cell nucleosome, methylation and transcription sequencing (scNMT-seq) provide us collective overview of chromatin accessibility, methylome and transcriptome on same cells^3^.

Single-cell multiomics analysis broadens our perspective on cellular diversity which is important for cellular development. This diversity takes the shape of cell populations and subpopulations which have direct clinical implications for both diagnosis and treatment disease. Different distributions of cell subpopulations in tumour microenvironments, for instance, is a key component for the prediction of immunotherapy response and gene discovery^4–6^. Increasing quantity of spatial omics data emphasises the importance of single-cell multiomics analysis especially in cancer research^7^.

In order to handle the high complexity, various computational tools have been developed to integrate single-cell multiomics data and analyse cellular heterogeneity based on the integration^8–10^. These analyses can address many unanswered questions in biology and medicine, such as novel cell type detection or relationships between different cells^11^. However, computational analysis of single-cell multiomics data faces several major challenges. For instance, sparsity in single-cell multiomics data could lead a statistical model to a wrong distribution due to a lot of missing values. Moreover, the data may include biases introduced by different subjects or workflows, which obscure other crucial biological signals showing cellular heterogeneity.

A variational autoencoder (VAE) is a deep learning model for mapping large input data to a latent space representation with reduced dimensionality^12^. It has been used for integrative analysis of multiomics data with applications in cancer classification and subtyping^13^. With the growing availability of single-cell multiomics data, VAEs have been used to summarise extremely complex and sparse datasets using interpretable latent factors^9,14,15^. Nevertheless, most VAE-based methods have only been tested on specific combinations of assays^9^, or do not properly address the sources of variation from different factors and batch effects^13,14^, which are the most critical issue in single-cell multiomics integration.

Here, we present a new single-cell multiomics integration method, Single-cell Multiomics Autoencoder Integration (scMaui). scMaui uses VAEs to integrate highly complex single-cell multiomics data into a lower dimension of latent space (Figure 1). scMaui particularly corrects batch effects using covariates and adversarial learning. The newly created latent space not only enables enhanced cellular heterogeneity analysis within thousands of single-cells, but also reveals hidden relations between the different modalities. Since VAE is an unsupervised learning method, users do not need to provide any reference data which is rarely available in single-cell multiomics analysis. Furthermore, scMaui can handle multiple batch effects independently via a direct conditioning on covariates, as well as using adversarial learning. We made scMaui robust to sparse data so that a missing assay in a subset of samples can be handled.

**Figure 1.**
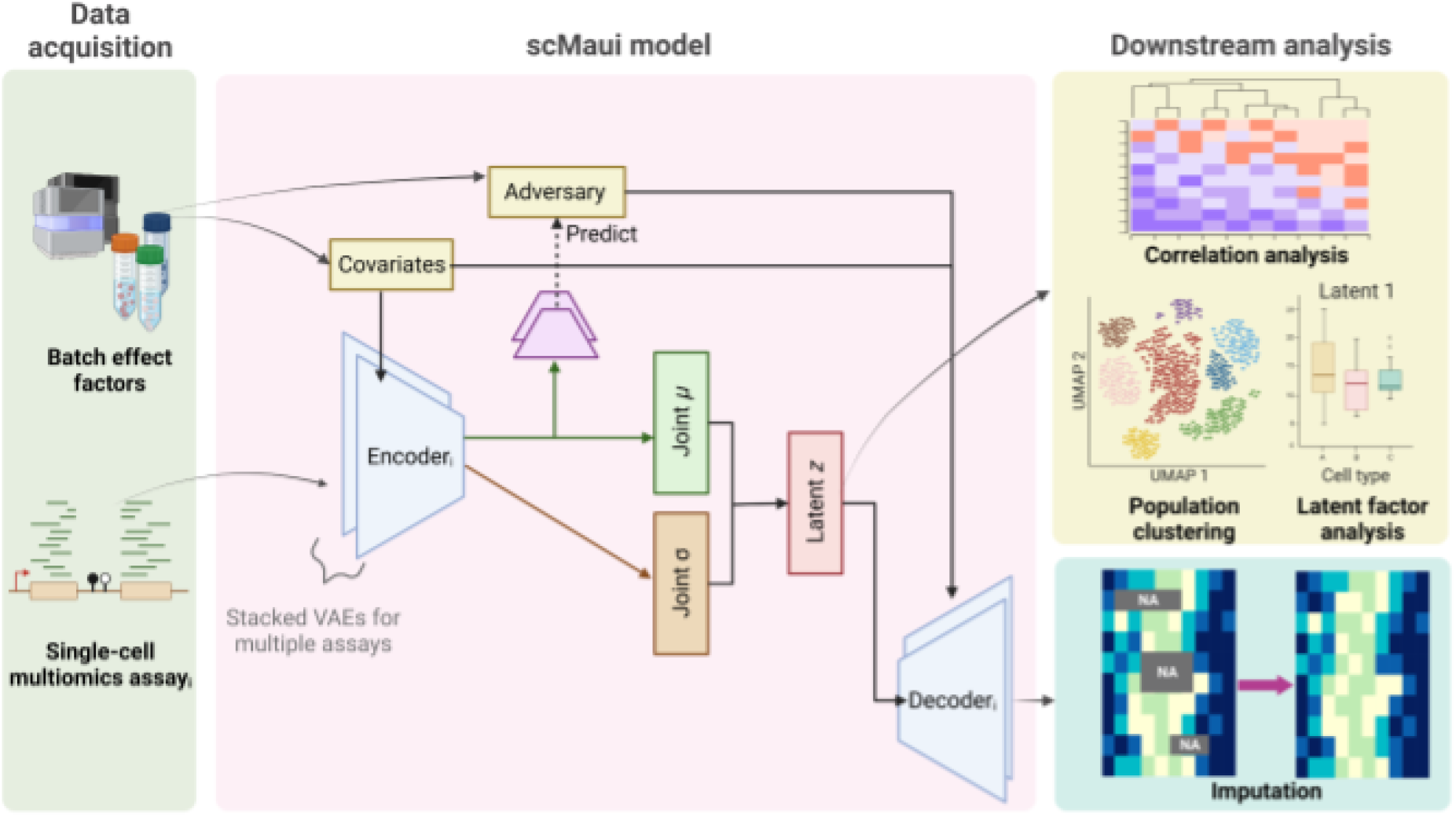
Illustration of scMaui model overview and the training process. Each single-cell multiomics assay is given to an encoder, whereas batch effect factors are independently handled by covariates and adversary networks. Latent factors created by scMaui can be used for downstream analyses to find cellular heterogeneity (e.g. sub/population clustering) and reconstructed assays by the decoders can be used for imputation.

We thoroughly assessed scMaui compared with other single-cell multiomics integration tools and conducted further biological analyses using various single-cell multiomics datasets including different assays. We show that scMaui outperforms other methods in many benchmarks, and is capable of cellular heterogeneity analysis across different biological samples. We also demonstrate the broad utility of scMaui through the analysis of diverse datasets and modalities.

## Results

### Evaluation of scMaui performance

Here, we compared scMaui to other previously published methods in terms of cell population and subpopulation identification. Each of the integration methods produces a reduced dimension latent space representation of the data. We used human bone marrow single-cell RNA-seq and antibody-derived tags (ADTs) dataset (GSE194122^16^) as our benchmarking dataset. We modified given cell-type labels within the dataset to *population* and *subpopulation*. Criteria to assign the labels are explained in Methods. For the evaluations, we used 84,677 cells in total including 11 populations which comprise 45 subpopulations. These are collected from 8 different donors and processed in 4 sites.

For the first benchmark, we sought to evaluate to which extent the cell type heterogeneity is preserved in the low dimensional representation (cell sub-/population prediction). scMaui created 50 latent factors in this task. To that end, we trained classifiers to predict the cell population from the latent space representation, and calculated the area under the receiver operating characteristic curve (AUC) to measure classification performance. We used a support vector machine with 10-fold cross validation and calculated average performance over all cell populations. Regarding average AUC, scMaui outperformed all alternative methods (Figure 2A). The performance difference was most striking for the innate lymphocyte cell (ILC) population as shown in Figure 2B. ILC population was particularly difficult to distinguish from T cells, because ILC and T cells share a lot of molecular signals and regulate several similar immune functions^17^. However, ILCs do not have an adaptive T cell receptor and belong to the innate immune system^18^, making them a distinct population which scMaui identified with better AUC than the other methods. We proceeded with the same analysis, but on cell subpopulation label ([Supplementary](https://docs.google.com/document/u/0/d/15y9HrqklP24RFPXez0am6lWFLZTkyyOGc4kFGi-EXBY/edit) Figure 1). MOFA, both with and without group option, outperformed other methods showing high performance in cDC1 subpopulation, but scMaui still achieved better mean AUC score than Seurat and totalVI.

**Figure 2.**
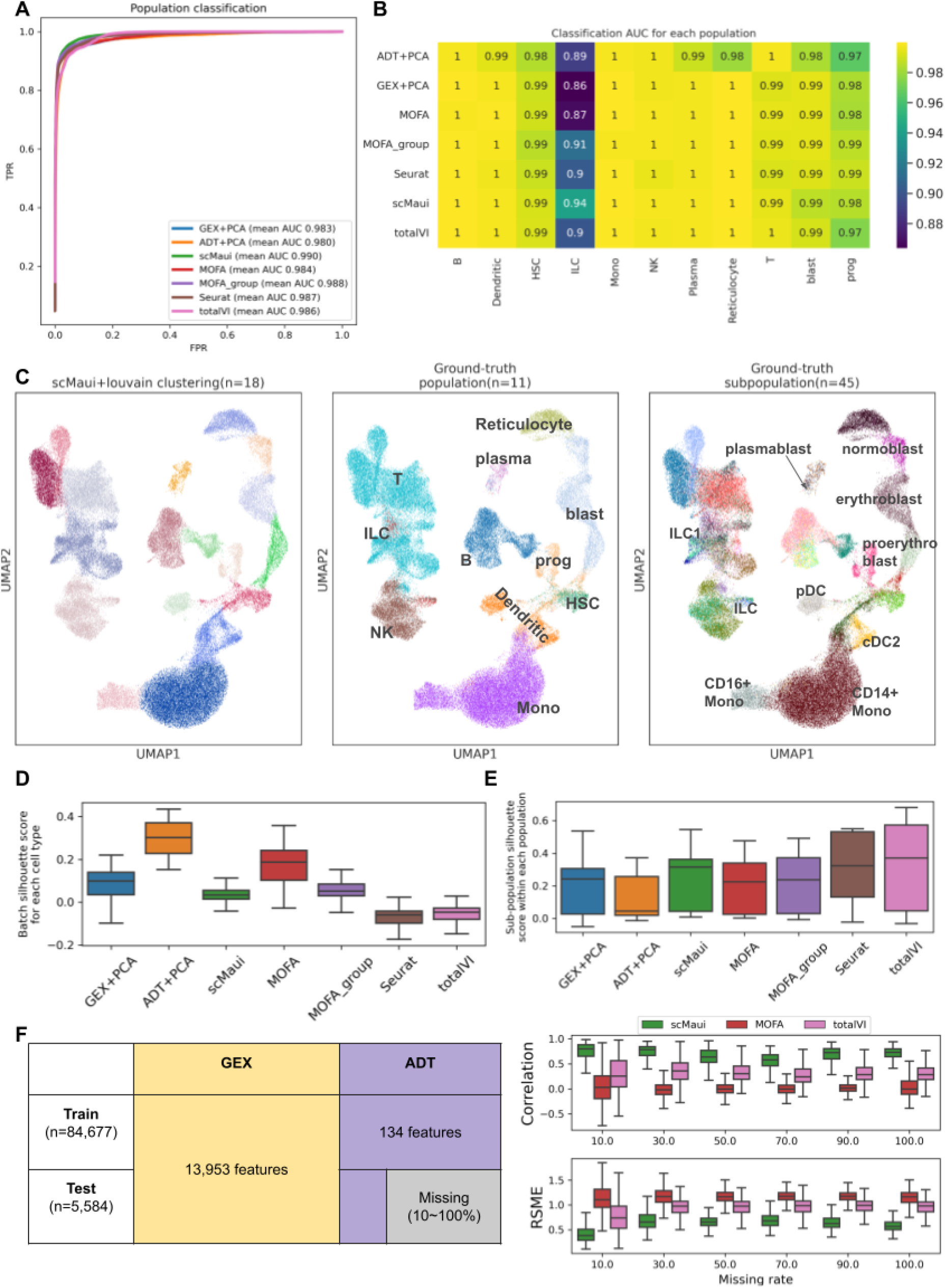
Benchmarking results of single-cell multiomics integration methods. **A.** Cell population classification receiver operating characteristic (ROC) curves and mean AUC. **B.** Classification AUC value for each population and each method. **C.** UMAP representation of scMaui latent factor coloured by clustering result, ground-truth population and subpopulation labels. **D.** Batch effect silhouette score in each subpopulation. **E.** Subpopulation silhouette score in each population. F. Protein expression (antibody-derived tags, ADT) modality imputation task dataset overview (left) and correlation results between predicted and ground-truth values. All boxplots present the median value as a middle bar in the box and both ends refer to as the first and the third quantiles.

In practice, ground-truth cell type labels are often not available for single-cell multiomics analysis. Therefore we also evaluated all methods in terms of clustering, rather than supervised classification. Louvain clustering was applied to the latent factors extracted by each method. After running Louvain clustering, we measured the clustering performance with adjusted mutual information (AMI), adjusted random index (ARI) and clustering purity (Table 1). To find the best clustering result for each method, 20 different resolution values (increased by 0.1 from 0.1 to 2.0) were tried and the value recording the best AMI score was selected. scMaui achieved the highest score in all these clustering performance measurements. Figure 2C shows the UMAP plot of scMaui latent factors with Louvain clustering results, ground-truth population and subpopulation respectively. In total, 18 clusters were detected via Louvain clustering algorithm, which falls into between the number of populations and subpopulations. The detected clusters covered most ground-truth populations and some subpopulations. In particular, reticulocytes, three subpopulations of blast and two monocyte subpopulations are covered by individual clusters very closely.

**Table 1.** Performance comparison with respect to clustering and dimensionality reduction. Louvain clustering algorithm was applied to the low-dimensional latent factors/features extracted by each method. Clustering results were assessed with AMI, ARI and clustering purity score, while population silhouette score indicates how well-separated populations are in the extracted latent factors/features. The best value of each score is highlighted in bold.

|  | Best resolution | AMI | ARI | Clustering purity | # detected clusters | Population Silhouette score |
| --- | --- | --- | --- | --- | --- | --- |
| GEX+PCA | 0.4 | 0.709 | 0.603 | 0.584 | 16 | 0.255 |
| ADT+PCA | 2.0 | 0.625 | 0.268 | 0.656 | 53 | 0.139 |
| scMaui | 0.7 | **0.796** | **0.737** | **0.722** | 18 | 0.274 |
| MOFA | 1.3 | 0.684 | 0.345 | 0.714 | 40 | 0.183 |
| MOFA_group | 0.4 | 0.753 | 0.703 | 0.658 | 18 | 0.157 |
| Seurat | 0.3 | 0.760 | 0.713 | 0.617 | 16 | **0.312** |
| totalVI | 0.9 | 0.773 | 0.542 | 0.710 | 25 | 0.296 |

The silhouette score measures how well-separated given sets of labels are within the feature space. We calculated the silhouette score for batch labels and population labels in the latent space created by each method. For batch labels, the score was computed within each subpopulation (Table 1 and Figure 2D). While high silhouette score is expected for well-separated populations, ideally batch-corrected latent space should record low silhouette score with respect to the batch labels. Regarding population and batch silhouette scores, Seurat performed best, but scMaui showed relatively high performance compared to MOFA and PCA applied to each assay. We furthermore calculated silhouette scores of subpopulations within each population in order to investigate how well-distributed subpopulations are in their population cluster (Figure 2E). scMaui achieved the third highest median subpopulation silhouette score following totalVI and Seurat.

We also assessed scMaui and other methods in a missing data imputation task (Figure 2F). Here, new test data was introduced from the same GEO database, but data from the protein expression modality was partially masked with different rates. For comparison, we included only MOFA and totalVI, which had a clear tutorial of protein expression imputation. scMaui imputed masked protein expression values most accurately showing the highest correlation and the lowest root mean squared error (RMSE) with respect to the ground-truth protein expression value, regardless of the missing rate.

### scMaui is capable of trajectory inference and subpopulation examination in human bone marrow samples

Subpopulation examination and trajectory inference (e.g. for normal or diseased cell differentiation and maturation) are among the most frequent use cases of single-cell multiomics analysis^19–21^. Cell populations and subpopulations can be annotated based on marker gene expression referring to already published single-cell atlases, but inconsistent definition of cell states and subpopulations between different reference datasets make it challenging to interpret single-cell multiomics data^22^. Another major challenge comes from the cells in transition between states. Transcriptomic signal change is continuous rather than discrete and cell fates are decided stochastically^23^. These alterations need to be thoroughly demonstrated for further analyses such as trajectory inference, which is also known as pseudo-time ordering and tracks the dynamic cell differentiation among single cells. Therefore, single-cell multiomics integration tools should be able to deal with the change of expression levels accordingly and extract features which can explain it based on multiple assays. We validated the usability of scMaui in these regards using single-cell RNA-seq and assay for transposase-accessible chromatin sequence (ATAC-seq) from healthy bone marrow samples (GSE194122 ^16^). Here, we used 63,138 cells provided by 9 donors and processed in 4 different sites. 10 populations including 22 subpopulations were given in the dataset. scMaui was set up to generate 50 latent factors as the previous task.

Firstly, we calculated the correlation between latent value and some T cell marker gene expression (Figure 3A). We selected *CD3D* as a marker for general T cells, *GNLY, CCL5, NKG7* as markers for CD8+ T cells, and *CD4*, *SELL, CCR7* as makers for CD4+ cells based on previous studies^10,16^. These three groups of markers resulted in the same three clusters based on the correlation, via hierarchical clustering, and the three CD8+ T cell markers explicitly showed similar patterns of correlation values. Moreover, CD8+ cells had higher values for the latent factor 38 and 36 (Figure 3B) which are also highly correlated with the CD8+ T cell marker genes. On the other hand, CD4+ T cells showed higher values than CD8+ T cells at latent factor 32 and 16, which the CD4+ T cell markers are highly correlated with. These results demonstrate that scMaui is capable of summarising cell-type-specific signals into its latent factors without prior knowledge about markers or cell-types.

**Figure 3.**
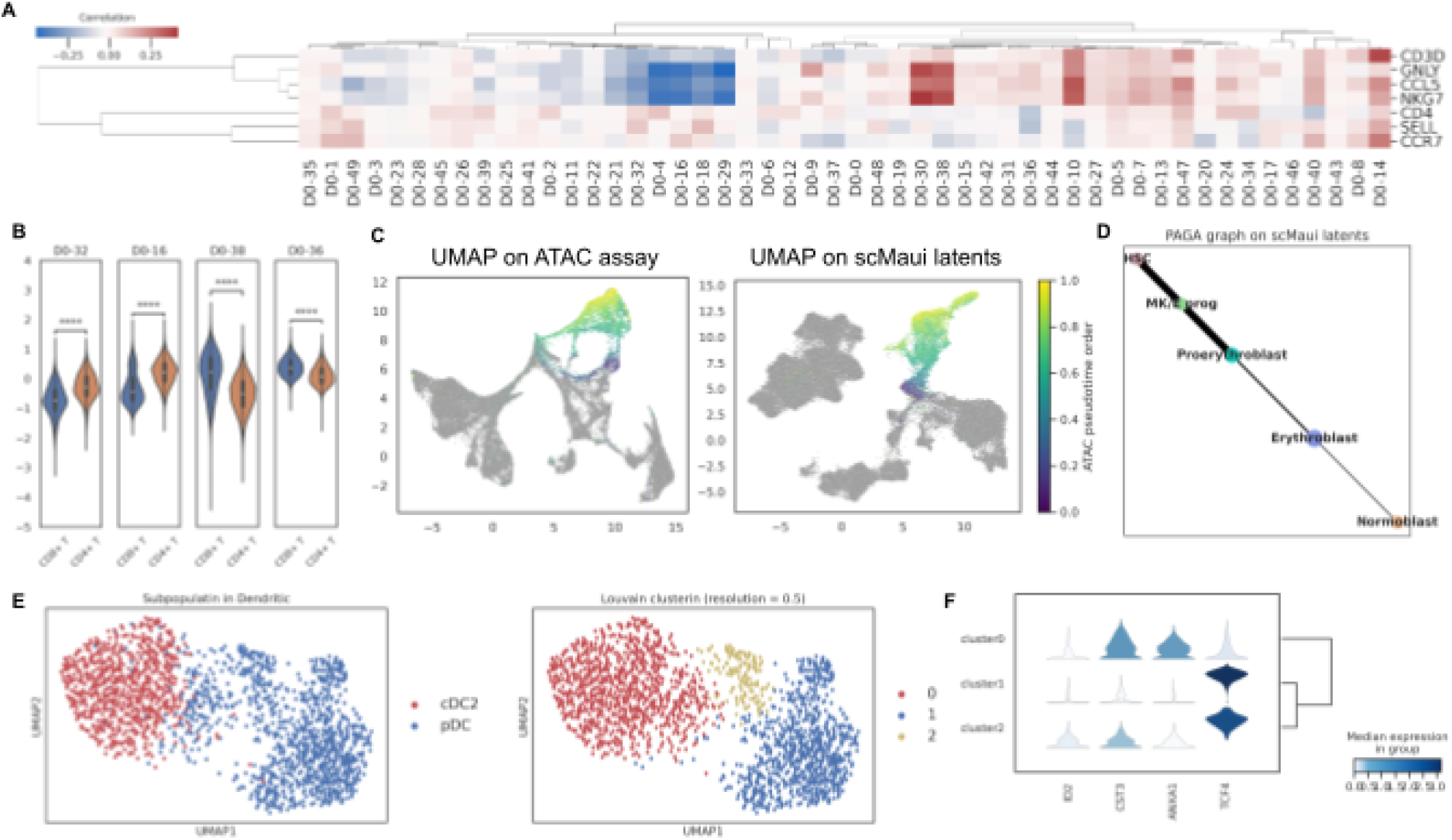
Subpopulation examination and cell-trajectory analyses using scMaui **A.** Correlation between 50 scMaui latent factors and T cell marker genes **B.** Distribution of latent values over CD4+ and CD8+ T cells. Only latent factors highly correlated with their marker genes were chosen. **C.** UMAP plot of PCA derived from ATAC-seq assay (left) and scMaui latent factors (right) coloured by ATAC-seq pseudo-time order **D.** PAGA graph applied to scMaui latent factor of HSC, MK/E progenitor and blast subpopulations **E.** UMAP plot of scMaui latent factor coloured by dendritic subpopulations (left) and louvain clusters (right) **F.** Dendritic subpopulation marker gene expression analysis in the detected louvain clusters

Figure 3C displays ATAC pseudo-time of HSC, MK/E prog and blast cells on UMAP plots calculated on principal components (PCs) of ATAC-seq assay and on scMaui latent factors. In this dataset, pseudo-time order of cells was generated in each modality using diffusion pseudo-time^24^. Compared to the UMAP computed directly from PCs, scMaui latent factors bettered the distribution of single-cell data accordingly with ATAC pseudo-time order. A partition-based graph abstraction (PAGA) map generated from scMaui latent factors also reflects the order of cell development from HSC to normoblast^25^ as a consistent result with the pseudo-time order (Figure 3D). PAGA was developed to graph a topological structure in single-cell data points based on partition connectivity, and has been used for precise identification of biological trajectories^26^. We performed the same cell-trajectory analysis with MOFA and Seurat ([Supplementary](https://docs.google.com/document/u/0/d/15y9HrqklP24RFPXez0am6lWFLZTkyyOGc4kFGi-EXBY/edit) Figure 2). While Seurat made a consistent result with scMaui, MOFA factors could not distinguish HSC, proerythroblast and normoblast as clearly as scMaui did and created weak connections between HSC and two blast subpopulations.

In further analysis, scMaui detected three clusters (using Louvain clustering as described above) in pDC cells, which are a subpopulation of dendritic cells (Figure 3E). To explore the discrepancy between the detected two clusters in pDC, we investigated expression level of some dendritic marker genes in all three clusters found in the dendritic population (Figure 3F). Cluster 0 mostly belongs to cDC2 cells according to both UMAP plots and expression analysis. However, cluster 1 and 2 show a difference mainly in ID2 and CST3 genes, which are considered as cDC1 cell markers rather than pDC cell markers according to Lin et al. and Schlitzer et al^27,28^. ID2 and CST3 genes are generally hyper-expressed in cDC1 cells. Therefore we presume that pDC cells in this dataset might include cDC1-like cells in the cluster 2, although the dataset does not contain cDC1 in their cell-type labels. This provides further evidence of scMaui’s superior ability to differentiate between subtle cell subtypes without prior knowledge.

### scMaui can explain both mouse embryo development and cellular heterogeneity based on DNA methylation and gene expression

DNA methylation, especially occuring at CpG sites, involves cell-type-specific signals as well as carries tumour heterogeneity in mammals^29–31^, and many computational methods have been published for such analysis based on bisulfite sequencing (BS-seq) data^32^. In particular, hypermethylation at gene promoters commonly behaves as a gene suppressor and can lead to dysregulated gene expression associated with tumour^33,34^. However, data sparsity is particularly severe in single-cell methylation compared to other assays and can bias statistical models. In addition, methylation is also relevant to gender, age and environmental influences. Thus, single-cell multiomics integration should ideally be able to distinguish signatures derived from different factors, so that the desired variation under the relevant conditions for a study do not get mixed with technical and other biological sources of variation.

During the development of the mouse embryo, both transcriptomic and epigenetic landscapes differ in each stage and cell differentiation^35^. Hence, we analysed mouse embryo development based on both assays using scMaui. The single-cell multiomics dataset was collected from GSE121708^35^. After removing cells based on quality control (QC) results, 939 cells from 8 cell populations were used in our analyses. The cells were acquired from 28 different embryos in 4 different stages. When the RNA-seq assay was analysed with PCA, strong batch effects were shown as epiblast and primitive streak cells tend to be clustered by samples rather than cell populations ([Supplementary](https://docs.google.com/document/u/0/d/15y9HrqklP24RFPXez0am6lWFLZTkyyOGc4kFGi-EXBY/edit) Figure 3A and C). The UMAP of PCs coloured by embryo stage also poorly recapitulates the change of embryo stage ([Supplementary](https://docs.google.com/document/u/0/d/15y9HrqklP24RFPXez0am6lWFLZTkyyOGc4kFGi-EXBY/edit) Figure 3B).

On the other hand, scMaui could alleviate these issues and made a more meaningful organisation of cell populations (Figure 4A right). scMaui created a latent space with 28 factors and could precisely distribute mouse embryo single-cells according to biological cell lineages ^35–37^ in Figure 4C. We also found that scMaui latent factors could be used to detect embryo developmental stages (4.5, 5.5, 6.5 and 7.5) within respective cell populations (Figure 4A left). For instance, epiblast population includes all four different stages, and the cells were aligned along the UMAP2 axis accordingly. We did the same analyses with MOFA and Seurat ([Supplementary](https://docs.google.com/document/u/0/d/15y9HrqklP24RFPXez0am6lWFLZTkyyOGc4kFGi-EXBY/edit) Figure 5) but both could not make a clear representation of embryo development stage changes as scMaui did. MOFA factors clearly distinguished between different stages but did not organise the clusters along the stage change. Seurat PCs could better arrange the cells according to the stage, but E4.5 cells are distributed together with other cells and not very distinct.

**Figure 4.**
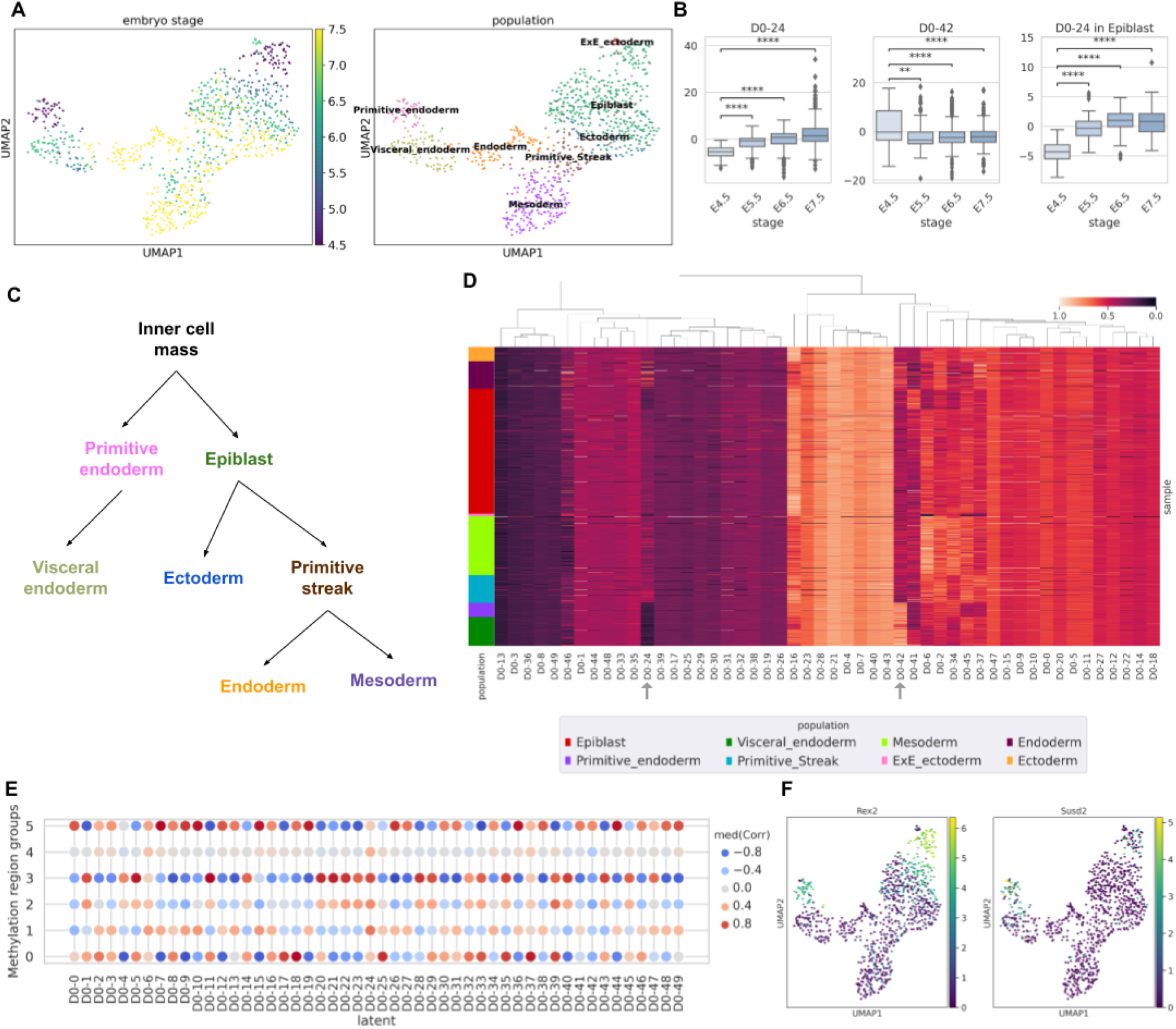
Mouse embryo single-cell gene expression and methylation multiomics data analysis results. **A.** UMAP plot of scMaui latent factor extracted from the entire dataset. The plots are coloured by embryo stages and populations each. **B.** Latent values in different stages of embryo cells. Latent factor 24 and 42 are presented at the left and the middle. The right graph shows the latent factor 24 values by embryo stage only in epiblast cells. **C.** Biologically proven embryo cell lineage. **D.** Latent values normalised between 0 and 1 and ordered by population. **E.** Median correlation between each latent factor and methylation level of each region group. The groups were decided based on clustering methylation levels. **F.** Gene expression of Rex2 and Sppr2a1 over all single-cell samples.

Among the fifty latent factors extracted by scMaui, latent factor 24 explicitly reflects the mouse embryo development stages (Figure 4B left). Although some epiblast cells present lower value than others in latent factor 24 (Figure 4D), there is still a clear increase of the latent value from stage E4.5 to E5.5, and from E5.5 to E6.5 (Figure 4B right). Latent factor 42 shows much higher values in the stage E4.5 cells than in the other stages of cells (Figure 4B middle) and, and primitive and some visceral endoderm cells, which are in E4.5 stage, also present very high values in latent factor 42 (Figure 4D).

In order to evaluate whether scMaui can detect methylation signals according to cellular heterogeneity, we calculated correlation between all latent values and methylation beta values in the regions selected as explained in Methods. We clustered the methylation regions into six groups using agglomerative clustering ([Supplementary](https://docs.google.com/document/u/0/d/15y9HrqklP24RFPXez0am6lWFLZTkyyOGc4kFGi-EXBY/edit) Figure 6) and analysed genes assigned to the regions in group 2 and 3. These groups were chosen due to the strong positive correlation with latent factor 24 and negative correlation with latent factor 42 (Figure 4E). Figure 4F depicts the expression of the *Rex2* gene (assigned to chr4:147019856-147023856 region in group 2) and *Susd2* (assigned to chr:75642008-75646008 region in group 3). Both are highly expressed in E4.5 cells (primitive endoderm and early epiblast populations). To sum up, these cells, where *Rex2* and *Susd2* are highly expressed, have high values in latent factor 42 and methylation level in promoter/enhancer regions of those present strong negative correlation with latent factor 42, which means these regions are hypomethylated. It corresponds to the main characteristic of CpG methylation in mammals that the genes are highly expressed when their promoter/enhancer regions are hypomethylated. Therefore, we claim that scMaui is capable of detecting a pair of relevant genes and methylation signals in terms of both mouse embryo development stages and cell-type heterogeneity.

## Discussion

Single-cell multiomics is growing in popularity due to the benefit of matched omics profiles from the same cells. However, the high complexity of single-cell multiomics data requires sophisticated computational tools to integrate assays into more comprehensive data representations, where cellular heterogeneity is clearly revealed and different sources of variation are disentangled. Different kinds of machine learning algorithms have been applied to this problem delivering better comprehension of molecular interactions between cells.

In this study, we presented scMaui, a stacked VAE-based single-cell multiomics integration model, and showed its capability of extracting essential features from extremely high-dimensional information in varied single-cell multiomics datasets. scMaui is trained to encode given data into a reduced dimensional latent space after processing each assay in parallel via separated encoders. During the training, it also adapts the latent space towards the direction that its decoder can reconstruct each assay as accurately as possible. In order to reflect the distribution of the different assays, scMaui supports multiple types of reconstruction errors. In addition, scMaui can handle multiple batch effect factors independently. Considering that different batch effect factors (e.g. patients, pipelines) introduce independent and irrelevant variations to data, alleviating different sources of variation separately in single-cell multiomics data as scMaui does, is statistically more coherent.

We assessed scMaui compared to several previously published methods: MOFA, Seurat and totalVI. Single-cell RNA-seq and protein expression data collected from human bone marrow samples were used for the evaluation. A summary of the evaluation is presented in Figure 5. scMaui outperformed other methods in population classification, population clustering and protein expression imputation tasks. Through the classification AUC values for each population, we showed that scMaui performs particularly well for cell populations sharing similar expression signals. For the cell population clustering analysis, scMaui recorded the best AMI, ARI and clustering purity scores. Finally, scMaui greatly outperformed all other methods in the imputation task, predicting much more accurate protein expression data regardless of how much of the data it was shown.

**Figure 5.**
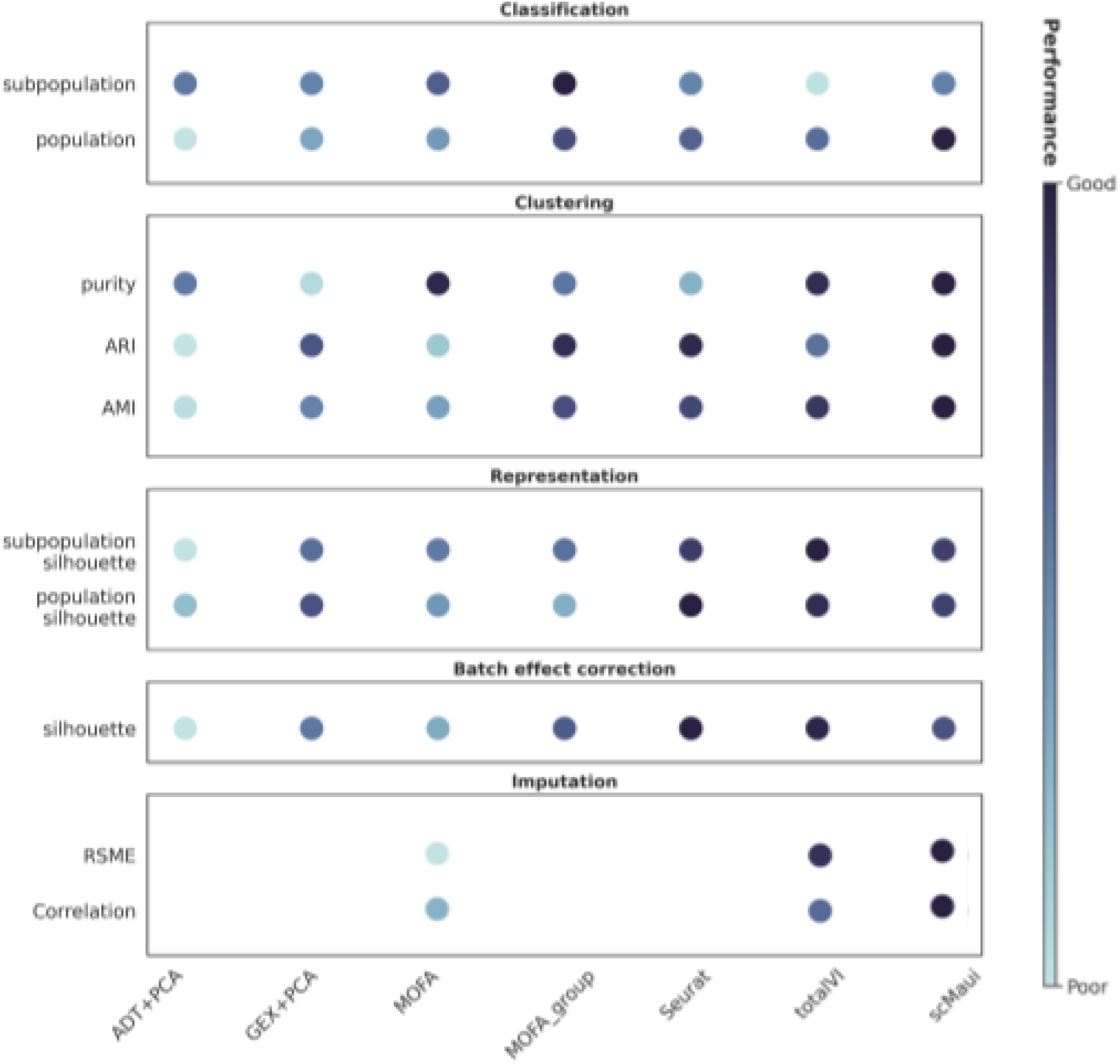
Summary of evaluation results conducted in our study. The darker the colour is, the better the method performed in each task. Performance scores are normalised divided by standard deviation in each task.

We conducted further downstream analyses of two more single-cell multiomics datasets with scMaui. Latent factors extracted from bone marrow single-cell RNA-seq and ATAC-seq dataset captured constant patterns of expression within CD4+ and CD8+ T cell marker gene groups respectively. Also, some of the latent factors which are highly correlated with marker genes presented higher values only in the corresponding cell type. These results indicate that scMaui can create its latent space corresponding to cell-type heterogeneity. scMaui latent space also showed a better organisation of HSC, MK/E progenitor and blast cells according to ATAC-seq pseudotime compared to MOFA factors and made a clear cell trajectory inference result. Lastly, based on the clustering result on scMaui latent factors, we found a new potential subpopulation in dendritic cells showing expression levels more similar to cDC1 cell type which is not annotated in the metadata.

DNA methylome provides a crucial perspective of epigenomic phenomena which induce different levels of gene expression. scMaui was able to create latent factors summarising relevant features in gene expression and methylation assays together, with single-cell mouse embryo multiomics dataset. Compared to other previous methods, the UMAP representation of scMaui latent factors arranged cells appropriately for both embryo development stages and cell populations. Also, we discovered that some scMaui latent factors can reflect the nature of general interactions between DNA methylation and gene expression, that hyper-methylation at CpG sites within promoter or enhancer regions impedes expression of the associated gene.

In conclusion, scMaui not only achieved better performance in many different analyses compared to other single-cell multiomics integration methods, but also showed its ability to reveal cellular heterogeneity and subpopulations remaining hidden in individual omics layers. From the technical standpoint, high flexibility supporting different types of reconstruction loss function, separated batch effect handling and missing feature indicator as described in Methods, are apparent benefits of scMaui pipeline. Thanks to this flexibility, scMaui covers more varied datasets produced via different workflows. Last but not least, scMaui provides a user-friendly and modern API relying on standard data structures. Consequently, we believe that the usability and the accessibility of scMaui will extend its usage to a broader range of researchers who would like to analyse their own data.

Although scMaui has shown outstanding performance and capability of cellular heterogeneity analysis, some questions still remain to be investigated. Because of the high cost of multi-assay measurements and numerous single-cell data already processed and stored in biobanks, there were attempts to integrate multiple layers of omics data collected from similar yet different cells^38,39^. This task mainly tries to create an integrated embedding space covering multiple assays and match clinically related cells between different assays. Since scMaui can process multiple modalities simultaneously and integrate those into a common embedding space, it might be possible to extend scMaui to multiomics data collected from unmatched cells or samples.

## Methods

### scMaui model

scMaui is composed of multiple VAEs as illustrated in Figure 1. One VAE network is assigned to each assay, so that assay-specific features can be extracted via separated networks. In order to create a joint embedding space for the integration, scMaui deploys additional layers calculating joint mean and joint standard deviation of encoded features from all assays. Joint standard deviation (*σ_joint_*) and joint mean (*μ_joint_*) for *N* assays are calculated as below:

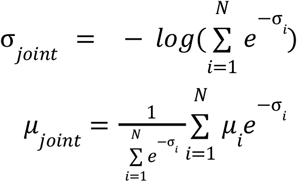

For joint standard deviation, we used the log-sum-exponential of standard deviation values computed from all encoders. In joint mean, we used a weighted sum of mean values that the smaller weight is given for the distribution with higher variance.

scMaui excludes sources of variations introduced by batch effects using two different techniques. Firstly, vectors storing batch effect factors are fed into encoders and decoders individually as covariates. Secondly, scMaui uses adversarial learning via additional feed-forward networks. The decoders reconstruct the input assays only from the jointly embedded latent factors and batch effect factor information conveyed by adversary and covariates.

Flexibility in batch effect correction is one of the strengths of scMaui. Batch effect factors can be both categorical and continuous values in scMaui model. Although varied statistical methods for batch effect correction have been proposed or included in multiomics integration pipelines ^40,41^, these are commonly limited to only categorical batch effect factors or even require more preprocessing such as separation of input dataset by batches. scMaui supports automatic and simultaneous correction of multiple batch effects ^42^.

Deep neural network models are optimised in a direction towards decreasing the given loss function value over multiple steps. scMaui has a loss function which consists of three terms:

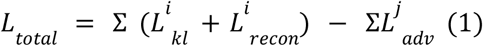

For each VAE assigned to an assay i, both KL-divergence (*L_kl_*) and reconstruction (*L_recon_*) losses are calculated as commonly done in VAEs training^43^. In scMaui, varied types of reconstruction loss function are supported depending on the assay type and the distribution of expression levels. For instance, negative binomial loss function was given to read count assays whereas methylation data was reconstructed based on binary loss for the bimodal distribution of methylation beta values. A full list of supported reconstruction losses is available in <u>Supplementary</u> Methods.

In addition, adversarial loss term (*L_adv_*) is included in the total loss function to correct batch effects within the input data. Adversarial learning ^44^ is a technique to force model training towards increasing certain loss term values on purpose. Since the latent space should ideally contain cellular heterogeneity information correcting all batch effects, the ground-truth batch effect vectors should not be predictable from the embedded latent factors. This idea is applied to scMaui loss function in a way of increasing cross-entropy loss between ground-truth and predicted batch effect values. According to the Equation 1, the model tries to decrease *L_total_* by increasing *L_adv_*. *L_adv_* is calculated for each batch effect factor *j* and summed up all together at the end. This approach was proven to improve batch effect correction by VAE-based models in our previous work^42^.

scMaui involves two unique benefits as a single-cell multiomics integration tool. Mosaic data where not all samples have complete modalities can be still modelled by scMaui using the “missingness” indicator. scMaui supports a “mask” layer where users can indicate which modality is absent for each sample, so that loss can be calculated only with measured assays and the robust embedding space can be created in spite of incomplete data.

Furthermore, different types of batch effect factors are handled in parallel as separate input vectors of covariates and adversaries. Continuous variables are also supported for batch effect handling in scMaui algorithm. These provide a high flexibility in scMaui downstream analysis for single-cell multiomics data, while users have to preprocess the batch effect information fitting to the required format (e.g. one category of all integrated batch effect factors) when using the majority of other single-cell multiomics integration tools.

### Data preprocessing

Three different single-cell multiomics datasets in various biological scenarios were used to assess scMaui performance. From GEO accession number GSE194122 ^16^, we downloaded two single-cell bone marrow datasets: one was comprised of RNA-seq and ADTs data, whereas the other includes RNA-seq and ATAC-seq. Mouse embryo single-cell multiomics data, which consists of RNA-seq and BS-seq, was also collected with GEO accession number GSE121708 ^35^.

Since each assay has its own properties, different preprocessing pipelines were applied accordingly. For RNA-seq and ATAC-seq assays, sequencing read counts were normalised and *log1p* transformed using scran ^45^ and scanpy^46^. Then, only for ATAC-seq assay, which is more sparse than RNA-seq, we excluded features covering less than 5% of the entire cells. ADTs were also normalised with the centred log ratio transformation.

Then, only for ATAC-seq and BS-seq assays, we filtered out features included neither in promoters nor in enhancer regions. hg38 and mm10 genomes were used for finding promoter and enhancer regions, as a human and mouse reference genome each. Promoters were collected from UCSC dataset, while enhancers were downloaded from FANTOM5 (https://slidebase.binf.ku.dk/human_enhancers/, ^47^). After the filtering, we selected only 5,000 most variable features from the assay.

Human bone marrow dataset (GSE194122) provides very detailed cell-type labels (45 labels for RNA-seq and ADTs data, 22 labels for RNA-seq and ATAC-seq data). Thus, we further coarsely annotated each cell by grouping given cell-type labels. In this study, the provided cell-type labels and the new group annotations are referred to as *subpopulation* and *population*. [Supplementary](https://docs.google.com/document/u/0/d/15y9HrqklP24RFPXez0am6lWFLZTkyyOGc4kFGi-EXBY/edit) Table 1 and 2 shows the annotation matches of population and subpopulation in individual dataset.

### Experimental setup

#### Assignment of batch effect factors

Depending on the single-cell multiomics data collection pipeline, different factors can introduce additional sources of variation within the dataset. For the human bone marrow dataset, two different factors were considered to cause batch effects: donor and site. On the other hand, in the mouse embryo development analysis, only different embryos were regarded as a batch effect factor. All the single-cell multiomics integration methods with the exception of scMaui took one merged vector of batches including both donor and site batch factors, because those accept only single type of factor for batch effect correction.

#### scMaui

Although scMaui requires many different hyperparameters regarding VAE setup, we kept consistent setup for most of those in all analyses. We gave 20 layers with 512 units to the encoder and 1 layer with 20 units to the decoder each. The adversary network was established with two layers having 128 units. Only in the input layer, dropout was applied with the rate 0.1. We used 512 as batch size in all model training.

Number of latent factors and training epochs were chosen depending on the number of features within the dataset in each analysis. While the human bone marrow gene expression and protein expression analyses used only 28 latent factors due to much lower number of protein expression features, other two analyses were done with scMaui trained for 50 latent factors. All models were trained over 1500 epochs with the exception of the human bone marrow RNA-seq and ATAC-seq analysis whose training was 2000 epochs owing to the larger number of features and less sparsity in ATAC-seq data compared to the methylation data. Regarding the reconstruction loss, we gave negative binomial loss to respective assays except the BS-seq assay in the mouse embryo development analysis. Since methylation data generally presents distinct binary signals, methylated or unmethylated, in single-cell data, we utilised binary reconstruction error.

#### PCA

Principal component analysis (PCA) is a linear dimensionality reduction algorithm which extracts the most variance components out of given data. We conducted PCA on each assay independently and extracted 20 components.

#### MOFA

MOFA ^8^ is an R package which can dissect multiomics dataset into integrated factors and provide a low dimensional reconstruction. In this study, MOFA2 version 1.6.0 was used and hyperparameters were chosen based on *get_default_model_options*, *get_default_data_options* and *get_defalut_training_options* functions provided by the package. MOFA also has an option to explain variations across different groups of samples, thus we did two separated analyses, with and without the group option, to compare the results in terms of batch effect correction.

#### Seurat

Seurat v3 ^10^ was mainly developed for integrating single-cell datasets generated via different experiments, but we used it in this study focusing more on its capacity to find associations between different assays. We applied *sctransform* only to the RNA-seq assay in all analyses. In order to handle the batch effect, the Seurat object was splitted into different batches. Then, we selected integration features with 3000 features, and found integration anchors between different assays. For finding integration anchors, we assigned 100 to the minimum number of neighbours as filtering threshold, ‘rann’ (it refers to as fast nearest neighbour search) to the neighbouring method. Ultimately, we ran PCA on the integrated data and extracted 20 components in each analysis.

#### totalVI

totalVI ^9^ is another VAE-based method for integrating specifically CITE-seq data, which is comprised of transcriptomes and epitopes, so it was only tested with the human bone marrow RNA-seq and ADTs data. We used totalVI included in python package scvi version 0.6.8. Batch label was given as the combination of doner and site. We set up batch size as 256, the ratio of training samples as 0.8 and the learning rate as 4 × 10 ^3^. The training was done over 300 epochs, and afterwards, we ran the posterior modelling with batch size 32. Otherwise, we followed up all default hyperparameter set-up described in https://docs.scvi-tools.org/en/0.6.6/index.html.

### Performance measures

#### Louvain clustering

Louvain community detection algorithm identifies clusters based on the relative density of connections between the inside and the outside of each cluster^48^. This method was applied to a broad range of single-cell analyses^49^. We clustered latent factors with the Louvain clustering algorithm and assessed the performance of respective methods.

#### Adjusted mutual information

Mutual information (MI) measures the similarity between ground-truth clusters and predicted clusters. Given *n* ground-truth clusters *U* = {*U*_1_, *U*_2_, …, *U_n_*} and *m* predicted clusters *V* = {*V*_1_, … *V_m_*}, MI between two clusters is given as:

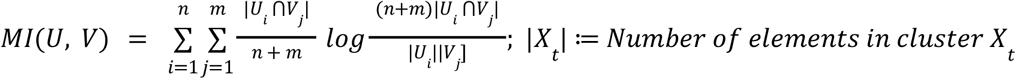

However, MI value tends to be higher when the number of clusters is larger. Therefore, we used adjusted mutual information (AMI) in our evaluation, which takes account of chance as follows:

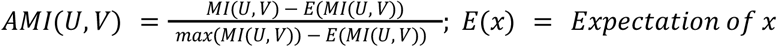

#### Adjusted rand index

Rand index (RI) is a statistics measuring the similarity between two different clusters. In our analysis, it was used for comparing predicted clusters and ground-truth clusters. RI is calculated as follows:

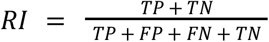

where *TP, TN, FP, FN* refer to true positive, true negative, false positive and false negative each. Adjusted rand index (ARI) is the corrected version of RI for chance and makes the measurement independent from the number of clusters. ARI equation is defined as:

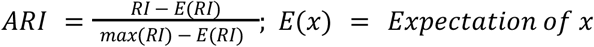

#### Clustering purity

We also calculated the purity of the most dominant class in each cluster. When *C* different ground-truth classes are grouped into one cluster, the clustering purity is calculated as follows:

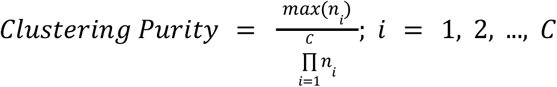

*n_i_* means the number of elements from the class *i*.

#### Silhouette score

For silhouette score values, we used silhouette coefficient which measures the similarity of each element to the cluster where it belongs to, compared to the closest neighbouring cluster. In single-cell analysis, it often represents the separation of clusters meaning that higher value indicates better separation. For each cell *c_i_*, silhouette score is calculated as below:

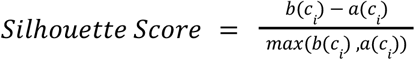

where *b*(*c_i_*) refers to the mean distance between the given cell *c_i_* and the nearest neighbouring cluster, on the other hand, *a*(*c_i_*) means the mean distance between the given cell *c_i_* and all other cells within the same cluster.

## Data availability

Both human bone marrow and mouse embryo single-cell multiomics datasets are available on GEO data repository with GEO accession number GSE194122 (ncbi.nlm.nih.gov/geo/query/acc.cgi?acc=GSE194122) and GSE121708 (ncbi.nlm.nih.gov/geo/query/acc.cgi?acc=GSE121708) each.

## Code availability

scMaui code is available on our Github repository (https://github.com/BIMSBbioinfo/scmaui). The github page also provides command-line usage and tutorials.

## Acknowledgement

We acknowledge Etienne Sollier's help (DKFZ) with proof-reading the manuscript.

