## Supplementary text, figures and tables for "Decoding single-cell multiomics: scMaui - A deep learning framework for uncovering cellular heterogeneity in presence of batch Effects and missing data"

### Supplementary Information

#### Methods

##### Reconstruction loss functions in scMaui

Here, we assume that the input assay $Y$ is comprised of $N$ features, $Y = [y_1, y_2, ..., y_N]$, and accordingly, scMaui decoder outputs the reconstructed assay $\hat{Y} = [\hat{y}_1, \hat{y}_2, ..., \hat{y}_N]$ with $N$ features. In the loss functions using binomial distribution, we specifically applied sigmoid function $S(x)$ to the output logit of the decoder as an activation function. Sigmoid function is defined as:

$$S(x) = 1/(1 + e^{-x})$$

**Poisson loss**

Poisson distribution, which originally models the probability that an event occurs the given number of times, is broadly used for omics data based on read counts. scMaui uses the negative log-likelihood of poisson distribution to calculate the poisson reconstruction loss as follows:

$$L_{poisson} = -\frac{1}{N}\sum_{i=1}^{N}(y_i log(\hat{y}_i) - \hat{y}_i)$$

**Negative binomial and negative multinomial distribution losses**

Due to the greater value of variance than the mean value, negative binomial distribution is considered as a better model for overdispersion than poisson distribution. For this reason, it is commonly used for explaining overdispersion in gene expression count data[1,2]. Thus, scMaui supports the negative log-likelihood of negative binomial distribution with parameters $r$ and $p$, as a reconstruction loss:

$$L_{negbinom} = -\frac{1}{N}\sum_{i=1}^{N}(log\Gamma(y_i + r) - log\Gamma(r) - log\Gamma(y_i + 1) + y_i log(S(\hat{y}_i)) + rlog(S(-\hat{y}_i)))$$

Since $p$, which refers to as probability of success in the binomial distribution, is assumed to be between 0 and 1 in negative binomial distribution, we converted the logit value outputted from the decoder with sigmoid function

On the other hand, in our previous work, negative multinomial distribution reconstruction loss outperformed negative binomial distribution loss with binarised single-cell ATAC-seq assay [3]. Therefore, we included it in scMaui package following the implementation in the previous work:

$$L_{negmul} = - log\Gamma(\sum_{i=1}^{N} y_i + r) + log\Gamma(r) - rlog(P_0) + \sum_{i=1}^{N} y_i \, log(P_i)$$

$P_0$ and $P_i$ are softmax-modified functions to make the non-negative parameters of multinomial distribution from the output of decoder $\hat{y}_i$. $P_i$ reflects each feature of input assay $y_i$ as follows:

$$P_i = \frac{exp(\hat{y}_i)}{1+\sum_{i=1}^{N} exp(\hat{y}_i)}$$

$P_0$ is used for explaining the dispersion in the data as below:

$$P_0 = \frac{1}{1+\sum_{i=1}^{N} exp(\hat{y}_i)}$$

**Binary loss**

Some single-cell omic assays, such as BS-seq or binarised ATAC-seq, resemble a bimodal distribution, so scMaui provides binary loss function to reconstruct this kind of assays. It is based on binomial distribution which models the number of successes in a sample size. The loss function again uses negative log likelihood of binomial distribution with a parameter $n$, the number of total trials, defined as:

$$L_{binary} = -\frac{1}{N}\sum_{i=1}^{N}( y_i log(S(\hat{y}_i))_- + (n_i - y_i)log(S(-\hat{y}_i)))$$

Due to the same reason as explained in negative binomial loss, we converted the output logit with sigmoid function.

**MSE and MAE**

Mean squared error (MSE) and mean absolute error (MAE) calculate error between the ground-truth and the reconstruction directly without any distribution:

$$L_{MSE} = \frac{1}{N}\sum_{i=1}^{N}(y_i - \hat{y}_i)^2$$

$$L_{MAE} = \frac{1}{N}\sum_{i=1}^{N}|y_i - \hat{y}_i|$$

| Subpopulation | Population | Subpopulation | Population |
|---|---|---|---|
| Naive CD20+ B IGKC+ | | ILC1 | ILC |
| Naive CD20+ B IGKC- | | ILC | |
| B1 B IGKC+ | B | Plasma cell IGKC+ | Plasma |
| B1 B IGKC- | | Plasma cell IGKC- | |
| Transitional B | | CD4+ T naive | |
| CD14+ Mono | Mono | CD4+ T activated | |
| CD16+ Mono | | CD4+ T activated integrinB7+ | |
| HSC | HSC | CD4+ T CD314+ CD45RA+ | |
| Reticulocyte | Reticulocyte | CD8+ T naive | |
| Normoblast | | CD8+ T CD49f+ | |
| Erythroblast | | CD8+ T TIGIT+ CD45RO+ | |
| Plasmablast IGKC+ | Blast | CD8+ T CD57+ CD45RA+ | |
| Plasmablast IGKC- | | CD8+ T CD69+ CD45RO+ | T |
| Proerythroblast | | CD8+ T TIGIT+ CD45RA+ | |
| NK CD158e1 | NK | CD8+ T CD69+ CD45RA+ | |
| NK | | CD8+ T naive CD127+ CD26- CD101- | |
| pDC | | CD8+ T CD57+ CD45RO+ | |
| cDC2 | Dendritic | MAIT | |
| cDC1 | | T reg | |
| Lymph prog | | gdT TCRVD2+ | |
| G/M prog | | gdT CD158b+ | |
| MK/E prog | Prog | dnT | |
| T prog cycling | | | |

***Supplementary Table 1.*** *Cell-type labels (subpopulation) and newly annotated population labels in GSE194122 single-cell gene and protein expression multiomics dataset*

| Subpopulation | Population | Subpopulation | Population |
|---|---|---|---|
| Naive CD20+ B | B | CD8+ T | T |
| B1 B | | CD8+ T Naive | |
| Transitional B | | CD4+ T naive | |
| CD14+ Mono | Mono | CD4+ T activated | |
| CD16+ Mono | | Lymph prog | Prog |
| Erythroblast | Blast | G/M prog | |
| Normoblast | | MK/E prog | |
| Proerythroblast | | ID2-hi myeloid prog | |
| NK | NK | pDC | Dendritic |
| ILC | ILC | cDC2 | |
| HSC | HSC | Plasma | Plasma |

***Supplementary Table 2.*** *Cell-type labels (subpopulation) and newly annotated population labels in GSE194122 single-cell gene expression and ATAC-seq multiomics dataset*

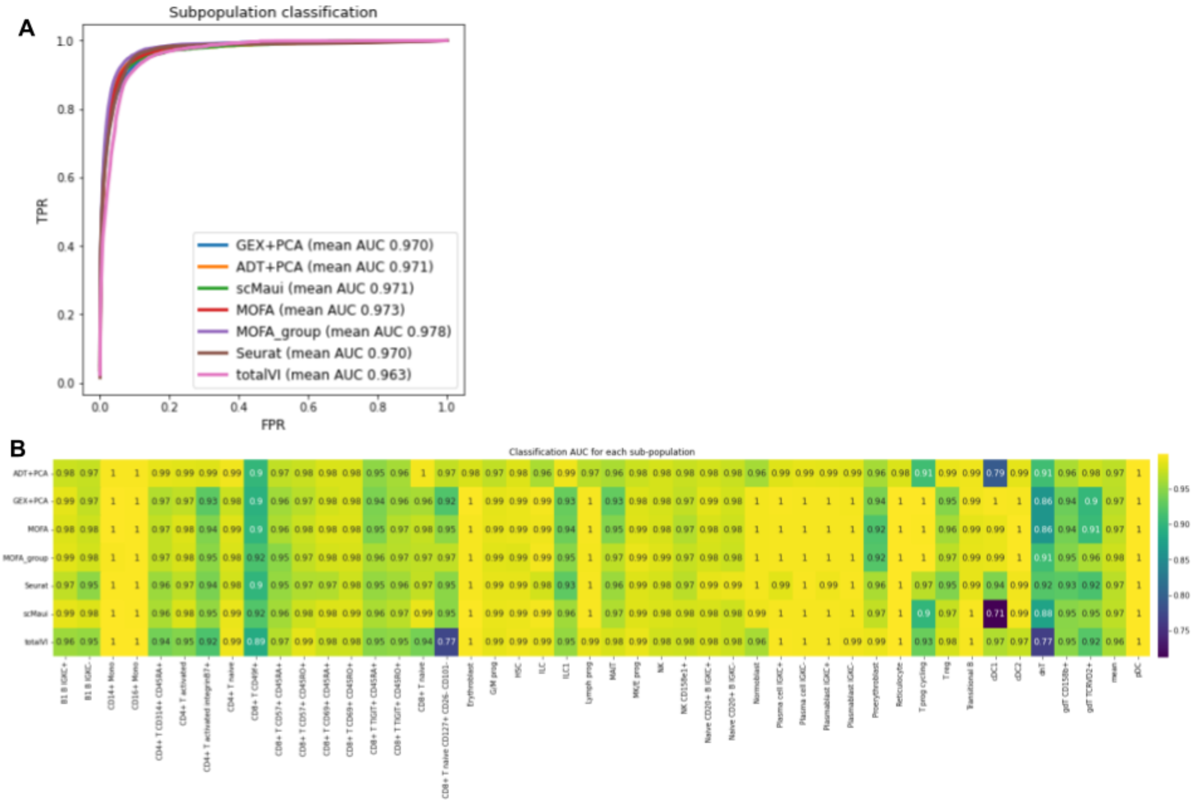

***Supplementary Figure 1.*** *Cell subpopulation classification results* ***A.*** *Cell subpopulation ROC curves and mean AUC.* ***B.*** *Classification AUC value for each subpopulation and each method.*

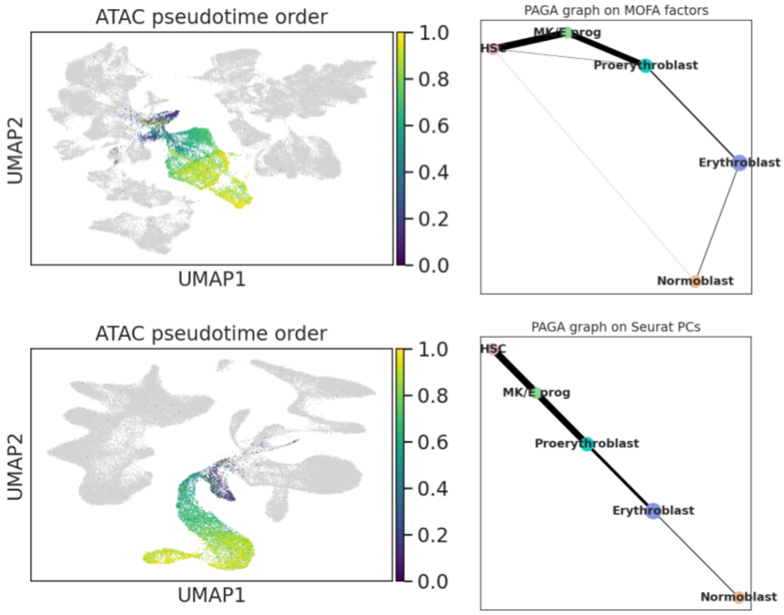


***Supplementary Figure 2.*** *ATAC pseudotime order representation on UMAP plots and inferred PAGA graphs of MOFA (top) and Seurat (bottom).*

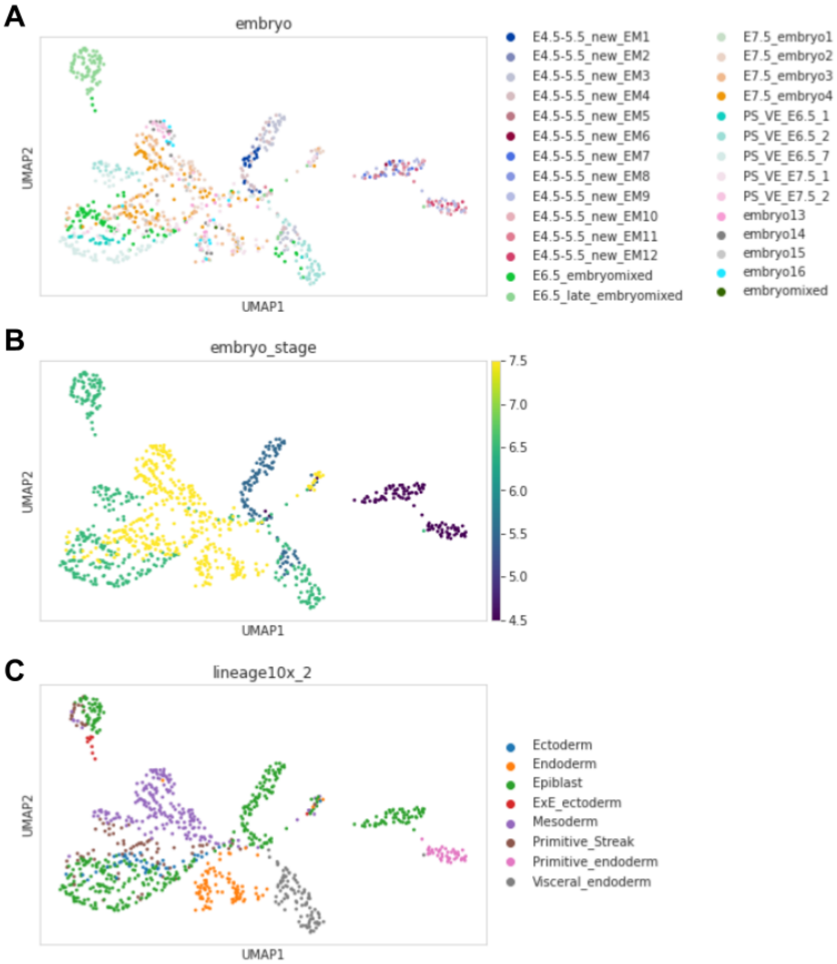

***Supplementary Figure 3.*** *UMAP plot of 20 principal components extracted from mouse embryo gene expression assay. A UMAP coloured by embryo samples B UMAP coloured by embryo stages C UMAP colored by cell-types*

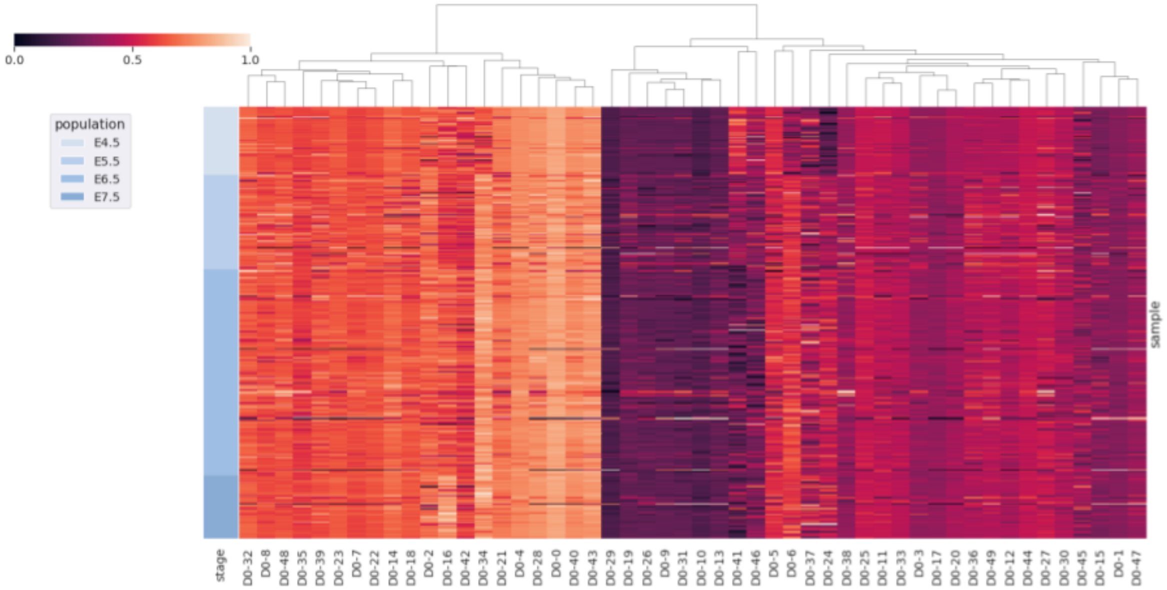


***Supplementary Figure 4.*** *scMaui latent values normalised between 0 and 1 and ordered by the embryo development stage.*

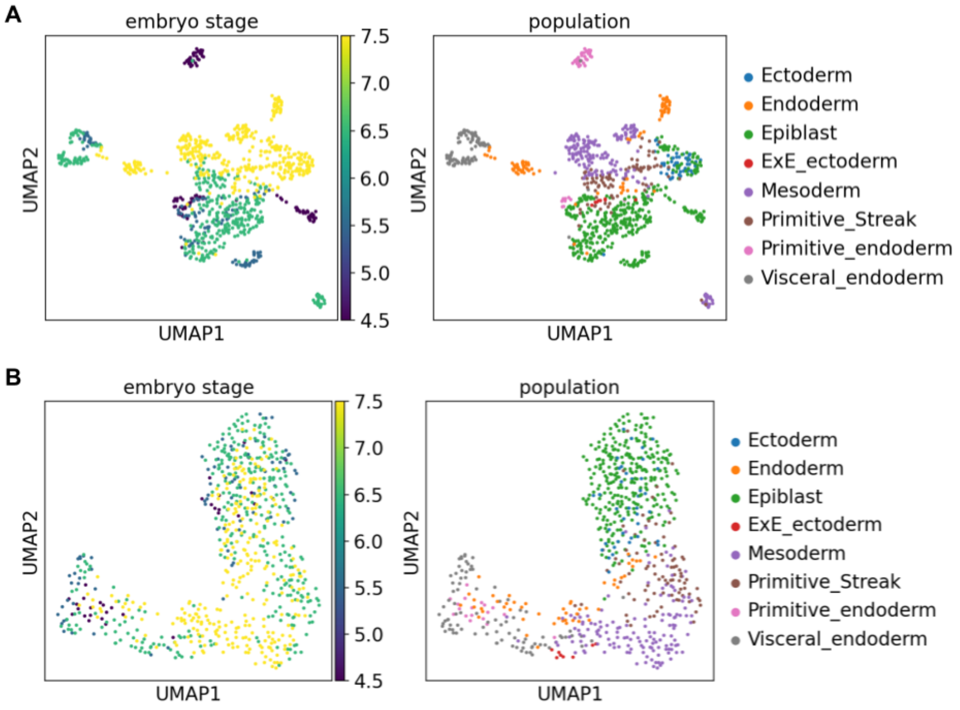


***Supplementary Figure 5.*** *UMAP plot of MOFA factors (A) and Seurat PCs (B) coloured by embryo stage and population*

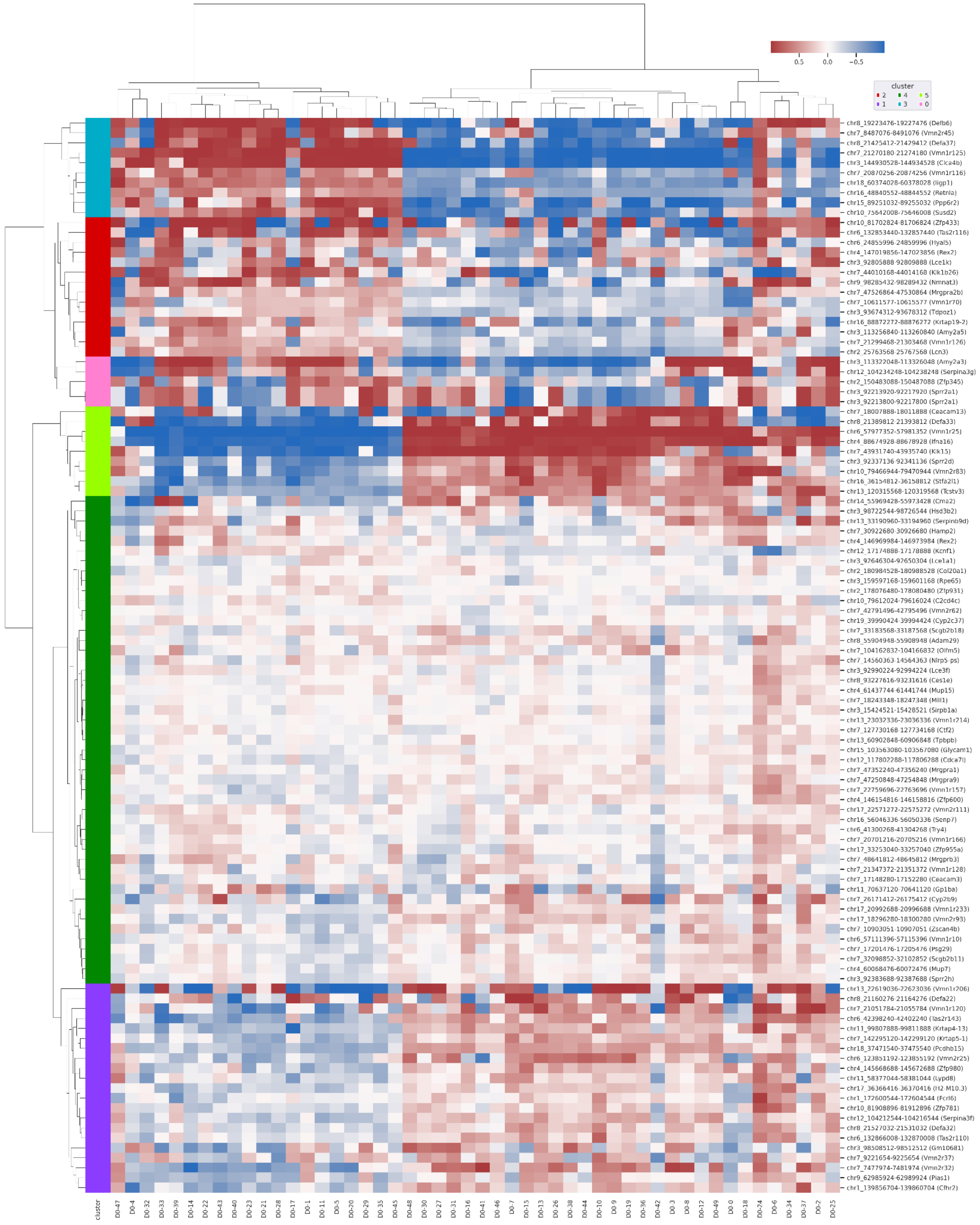


***Supplementary Figure 6.*** *Correlation between methylation level in promoter/enhancer regions and scMaui latent factors. Based on the correlation, we grouped regions into six clusters using agglomerative hierarchical clustering method.*
